# Detecting patterns of species diversification in the presence of both rate shifts and mass extinctions

**DOI:** 10.1101/004572

**Authors:** Sacha Laurent, Marc Robinson-Rechavi, Nicolas Salamin

**Affiliations:** Department of Ecology and Evolution, University of Lausanne, 1015 Lausanne, Switzerland; Swiss Institute of Bioinformatics, Quartier Sorge, 1015 Lausanne, Switzerland

**Keywords:** Phylogeny, Diversification, Mass Extinctions

## Abstract

Recent methodological advances are enabling better examination of speciation and extinction processes and patterns. A major open question is the origin of large discrepancies in species number between groups of the same age. Existing frameworks to model this diversity either focus on changes between lineages, neglecting global effects such as mass extinctions, or focus on changes over time which would affect all lineages. Yet it seems probable that both lineages differences and mass extinctions affect the same groups. Here we used simulations to test the performance of two widely used methods, under complex scenarios. We report good performances, although with a tendency to over-predict events when increasing the complexity of the scenario. Overall, we find that lineage shifts are better detected than mass extinctions. This work has significance for assessing the methods currently used for estimating changes in diversification using phylogenies and developing new tests.

## BACKGROUND

The estimation of the rates of speciation and extinction provides important information on the macro-evolutionary processes shaping biodiversity through time (Ricklefs 2007). Since the seminal paper by Nee *et al.* Nee et al. (1994), much work have been done to extend the applicability of the birth-death process, which now allows us to test a wide range of hypotheses on the dynamics of the diversification process.

Several approaches have been developed to identify the changes in rates of diversification occurring along a phylogenetic tree. Among them, we can distinguish between lineage-dependent, trait-dependent, time-dependent and diversity-dependent changes. Lineage specific methods identify changes in macro-evolutionary rates — speciation and extinction rates — at inner nodes of a phylogenetic tree (Rabosky et al. 2007; Alfaro et al. 2009; Silvestro et al. 2011). We can also identify trait-dependence in speciation and extinction rates — λ and *μ*, respectively — if the states of the particular trait of interest are known for the species under study (Maddison et al. 2007; FitzJohn et al. 2009; Mayrose et al. 2011). It is also possible to look for concerted changes in rates on independent branches of the phylogenetic tree by dividing the tree into time slices (Stadler 2011a). Finally, diversity-dependent effects can be detected when changes of diversification are correlated with overall species number (Etienne et al. 2012). Most methods can correct for incomplete taxon sampling, by assigning species numbers at tips of the phylogeny (Alfaro et al. 2009; Stadler and Bokma 2013), or by introducing a sampling parameter (Nee et al. 1994). By taking into account this sampling parameter at time points in the past, one can also look for events of mass extinction (Stadler 2011a).

These methods provide insights into the dynamics of species diversification, and it is now well accepted that differences in lineage-specific rates exist (Jetz et al. 2012; Barker et al. 2013). However, it seems unlikely that both lineage specific shifts and mass extinction events would not have occurred, especially when studying large phylogenetic trees covering hundreds of million years of evolution. For example, several global crises, which caused the extinction of a high proportion of species (Raup and Sepkoski 1982), have occurred since the appearance of the last common ancestor of vertebrates. Among them, the Cretaceaous-Paleogene (K-Pg) boundary and the Permian-Triassic events, which happened 65 million years ago (Mya) and 251 Mya, respectively, induced the most dramatic losses of biodiversity (Erwin 2006). Moreover, other less extensive events have also occurred in the past hundred million years (Benton 1995).

Mass extinction events could impact biodiversity in different ways. Three main hypotheses, corresponding to different patterns of extinction (Raup 1992), have been proposed. First, the event could affect all lineages equally and terminate any extant lineage with the same probability. This “field of bullets” scenario is often used as a null model (Nee 1997; Faller et al. 2008). Second, in the “fair game” scenario, some form of lineage selection would occur, where the most successful species — in our case, the most diversifying species — before the event would be the most likely to survive. This could, for instance, happen if the probability of survival depends on a specific trait varying across the lineages of the phylogeny(Faller and Steel 2012). Finally, in the “wanton destruction” scenario (Eble 1999), the event could induce such changes in the environmental conditions that the probability of extinction of the species and their post-event diversification rate would be uncorrelated to their initial speciation and extinction rates.

Although lineage-dependent differences in macro-evolutionary rates and mass extinctions are known to happen, the performances of the existing methods to identify both lineage-specific rate shifts when mass extinctions have occurred, and mass extinctions when lineage-specific rate shifts have occurred has not, to our knowledge, been investigated. The aim of this study was thus to assess the performance of current methods to estimate the rates of diversification using complex scenarios involving both mass extinctions and lineage shifts. We used simulations to assess the impact of varying number and magnitude of rate shifts and mass extinction events.

## METHODS

Figure 1 gives an overview of the simulation design. We used a backward algorithm to simulate phylogenetic trees as implemented in the *R* (R Core Team 2013) package TreeSim (Stadler 2011b), since a direct forward approach to simulate trees using a birth-death process can lead to bias when conditioning on the number of tips (Hartmann et al. 2010). The algorithm takes as input the number of extant species, the evolutionary rates λ and *μ*, and the time of occurrence and survival rate *ρ* for mass extinction events. We assume first that these events happen according to the field of bullet scenario (step 1). We randomly combine different trees having experienced the same mass extinction events but different evolutionary rates to account for rate shifts in diversification (step 2; see Table 1). This was done by ranking trees in decreasing order of their total age, which includes here the stem branch length provided by TreeSim. We selected from the oldest tree (referred to as acceptor tree) the branches that overlapped in time with the age of the stem branch of the second oldest tree (referred to as donor tree). Thus, the branches considered for possible grafting were the ones that included the age of the donor tree between the timing of the two speciation events defining them in the acceptor tree. We randomly chose one of those branches to graft the donor tree onto the acceptor. This ensures ultrametricity of the newly created tree, and leaves the branch lengths of each separate tree unmodified once the lineage having experienced the diversification shift is removed. We iterate over this protocol until all donor trees, whose number can vary between 0 and 5 (Table 1), have been grafted. Finally, we ran Medusa (Alfaro et al. 2009) and TreePar (Stadler 2011a) analyses on the resulting trees to investigate our capacity to recover the signal for mass extinctions and diversification shifts (step 3). We simulated trees with different numbers of lineages and of extinction events to account for their influence on the final results. Table 1 summarizes the parameter space explored for the 16371 trees that we simulated. For the values of λ and *μ*, we targeted distributions similar to the estimates calculated on a mammalian phylogeny (Bininda-Emonds et al. 2007).

**Table 1 –.**
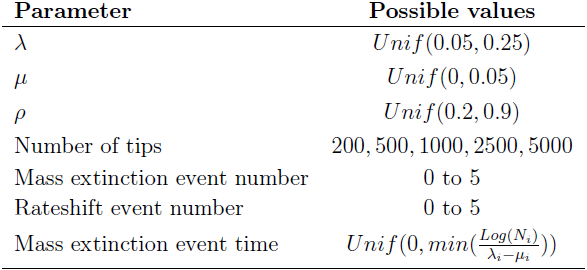
Universe explored for parameters values. *Unif*: Uniform distribution, *i*: lineage identifier

| Parameter | Possible values |
| --- | --- |
| $\lambda$ | $Unif(0.05, 0.25)$ |
| $\mu$ | $Unif(0, 0.05)$ |
| $\rho$ | $Unif(0.2, 0.9)$ |
| Number of tips | 200, 500, 1000, 2500, 5000 |
| Mass extinction event number | 0 to 5 |
| Rateshift event number | 0 to 5 |
| Mass extinction event time | $Unif(0, \min(\frac{Log(N_i)}{\lambda_i - \mu_i}))$ |

**Figure 1 –.**
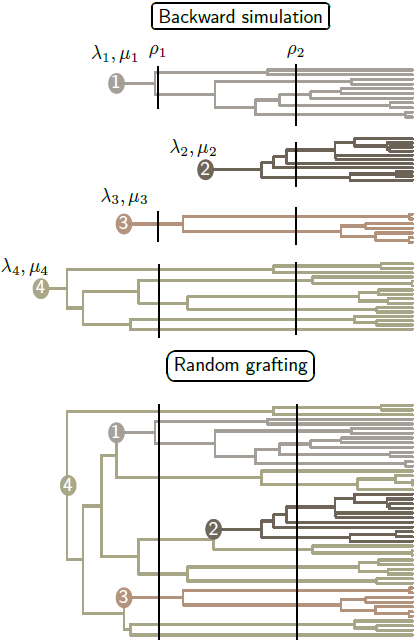
Workflow of the simulation process. Hypothetic case of 50 species tree, 3 lineages shifts and 2 mass extinctions. The number of species in each lineage is randomly drawn first. Each tree is grown separately with different (λ, *μ*) but with identical survival rates (*ρ*) at each mass extinction events. The four trees are then successively joined at branches ensuring ultrametricity. Vertical continuous lines: simulated mass extinction events, full circles: ancestor where diversification change occurred.

Medusa uses a maximum likelihood framework to detect shifts in diversification by iteratively adding breakpoints on inner branches of the tree with different rates of speciation and extinction. It uses AIC to discriminate between models with an increasing number of parameters (Alfaro et al. 2009). Medusa was run until a more complex model was not supported by the AIC. We did not extract the macro evolutionary rate estimations from Medusa as we were only interested in testing the ability of the method to detect the events, and not the accuracy of the parameter estimation.

TreePar uses the birth-death process to identify changes in λ and *μ* through time. This is done by estimating the probability of a change in parameter values within small time intervals, which can be extended to test for the occurrence of mass extinction events (Stadler 2011a). Given that the parameters of the rate shifts might be correlated with those related to mass extinction (Stadler 2011a), we restricted our analyses to the identification of mass extinction events. The number of iterations of TreePar was set to the simulated number of mass extinction events plus one to test for the appearance of false positive events. A standard Likelihood Ratio Test (LRT) is used to extract the most likely models from TreePar and more complex models were favored when their p-value was less than 0.01, following the standard approach for this framework (Stadler 2011a). Similarly to what was done with Medusa, we did not analyze estimations of survival rates at mass extinctions events given by this framework.

To verify that our simulation design has no influence on the methods we evaluated, we tested the influence of the subtree grafting approach with a constant rate of diversification. We simulated trees with 200 species using both the standard procedures implemented in TreeSim and by grafting two subtrees of 150 and 50 species having evolved under the same λ and *μ* values. We then compared the results obtained by TreePar and Medusa. We ran 250 pairs of simulations and we observed no significant differences in the number of false positive found between the groups with and without artificial grafting (7 and 13 for Meduse respectively, and none in both cases for TreePar), showing that our simulation design does not bias the estimation of the rate shifts by the two methods used.

We used a slightly different framework to study the impact of the different types of mass extinction events. We simulated a scenario that aimed at testing for the presence of the K-Pg mass extinction event using high order phylogenetic trees. We therefore simulated trees with a large number of extant species (5, 000 tips, similar to the number of mammalian species) and a large number of lineage shifts (5), but only one event of mass extinction. The other parameters were still drawn at random from the ranges specified in Table 1, except for the survival rate *ρ* that was modified according to the models of mass extinction. For the fair game hypothesis, we randomly drew λ and *μ* for the 5 different lineage shifts, but the survival rate *ρ* was modified for each lineage based on its diversification rate (λ – *μ*). We thus considered that the trait influencing the probability of extinction for each species was its diversification rate. For the wanton destruction hypothesis, the mass extinction event induced a change in rates for each lineage, again drawn according to the distribution stated in Table 1, and their survival rate *ρ* was then based on their new diversification value. For the wanton destruction, our simulations included both a global rate shift and a mass extinction and we ran TreePar twice in order to detect both events. For the two latter cases, we chose to linearly parameterize *ρ* with regards to diversification. As diversification could range between 0 and 0.25 and *ρ* between 0 and 1, we applied a factor four to the diversification to obtain the survival rates of the lineages. We also ran Medusa on the three sets of simulations to assess the potential impact of the three extinction hypotheses on the detection of lineage shifts. For this second part, we generated over 700 trees for each model of mass extinction event, for a total of 2289 simulations.

## RESULTS AND DISCUSSION

### Baseline Performances

To estimate the baseline behavior of both frameworks, we first tested the performance of the methods on the simplest scenarios. We thus selected simulations that included a single rate shift for Medusa, or a single mass extinction for TreePar. Figure 3 represents the fraction of shifts detected by Medusa relative to the absolute difference between the new and the old diversification values (λ – *μ*) (Figure 3A) and to the number of species in the lineage (Figure 3B). More than 80% of the changes in diversification larger than 0.05 were detected by Medusa, a good performance in assessing strong shifts. Further, Figure 3B shows that the overall tree size has no influence on the detection, since lineages of the same size are as likely to be detected in small or larger trees.

**Figure 2 –.**
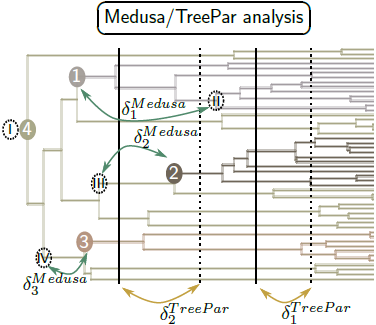
Exemple output of the analyzes. We run the Medusa and TreePar analysis, and group the pairs of simulated/estimated events by minimizing the sum of the distance separating the events in each pair 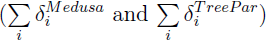. Vertical dotted lines: estimated mass extinction events by TreePar, dotted circles with roman letters: estimated diversification rate shift by Medusa, by decreasing significance, other: as in figure 1. The first estimated shift is always at the root of the tree.

**Figure 3 –.**
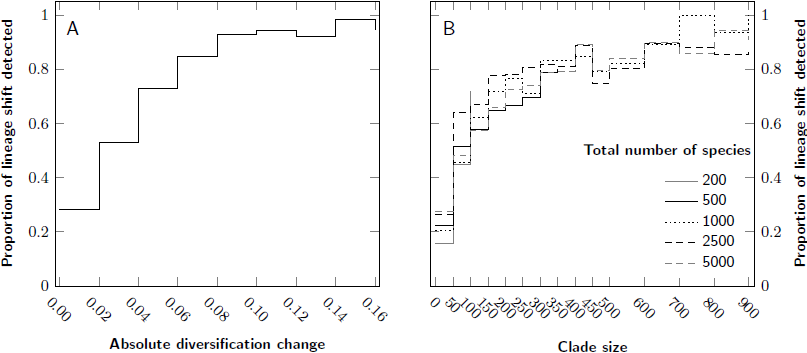
Baseline detection level for Medusa, for simulations with one rate shift and no mass extinction event. A: Proportion of detected events for ranges of values of diversification, B: Proportion of detected events for ranges of extant species number in lineages.

We then checked the ability of TreePar to detect the survival rate, *ρ*, associated with a mass extinction, as well as the number of ancestral species predating this event in the reconstructed tree. We also used first the simplest simulation to limit the effect of other parameters. Figure 4A shows that the signal of mass extinction in the phylogenetic tree is very weak when less than 100 ancestral species are present before the event. This has implications for our ability to find evidence for the K-Pg boundary using phylogenetic trees of vertebrates, for example. We can only reach more than a hundred ancestral species older than 65 My by considering phylogenetic trees encompassing distantly related lineages of tetrapods (see Bininda-Emonds et al. (2007) or Meredith et al. (2011)). Besides, as detection drops with increasing survival rate (Fig. 4B), the signal is even less likely to be picked as the ancestors of the extant species might have experienced the mildest extinction rates.

**Figure 4 –.**
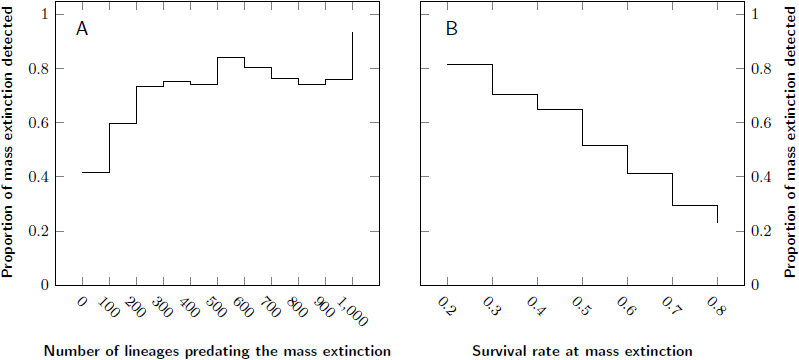
Baseline detection level for TreePar, for simulations with one mass extinction and no diversification shift. A: Number of lineages predating the mass extinction event influence, B: Survival rate influence.

### Mixed Scenarios of Diversification

In a second stage, we analyzed simulations with more events and a mix of different types of events. We evaluated the performance of rate shift detection by Medusa, or of mass extinction events by TreePar, by comparing the events detected to the relevant real (simulated) events. To perform the assignment between detected and simulated events (see Fig 2.), we chose to minimize the sum of the distances between each potential pairing of events 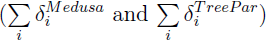. The distance metric used for Medusa is the sum of the branch lengths along the shortest path separating the two nodes, whereas we used the time between the estimated and simulated pairs of mass extinction events for TreePar (see caption of Figure 2 for details).

The simulations incorporate several factors and we tested the effect on the frameworks of three categorial parameters: total number of tips, number of mass extinctions and number of shifts in diversification rate (see Table 1 for their possible values). To ensure that the effects observed were related to the parameter of interest, we designed a reshuffling scheme for each parameter. First, we randomly selected an equal number of simulations for each combination of every possible value of the other two parameters. As an example, to study the outputs for trees of 200 tips, we randomly drew an equal number of simulations with (i) no lineage shift, no mass extinction and 200 tips; (ii) one lineage shift, no mass extinction and 200 tips; (iii) one lineage shift, one mass extinction and 200 tips; etc. This drawing was repeated a hundred times and we determined, for each bin created, the proportion of simulations for which each method favored the model with the correct number of relevant events it was looking for, and the proportion of simulations for which they favored a model with too many events. Finally, we report the median and 95% intervals of those proportions based on our hundred bins.

#### Tree size influence

Both Medusa and TreePar performed better in assessing the correct number of events they were set to detect with an increasing number of tips (Fig. 5). The median proportion of simulations correctly assessed reached 60% for Medusa and 32% for TreePar with 5,000 tips. The increase in the number of tips also led to an increased acceptance by TreePar of models with too many mass extinctions (28% for 5,000 tips). However, the number of tips in the tree has no effect on the error of the estimated time of mass extinction (Fig. 6), even though more events are predicted. We only see a slight effect of tree size for Medusa, which is probably due to the fact that the method only detects lineage related events and does not depend on the total number of tips. We also investigated the effect of lineage size on the outputs of Medusa. We first compared the variance of lineage sizes relative to the overall tree size, contrasting the simulations with false positives to those with the correct number of rate shifts found. To discard the effect of lineage number, we compared groups of trees with the same number of diversification shifts. To account for a potential effect of tree imbalance, we compared the variance in lineage sizes inside trees, with or without false positives. There was no effect in most cases, except in the simulations with 4 or 5 rate shifts (p-values: 0.01 and 3.6 · 10^−3^, respectively, Mann-Whitney test). Thus, simulations with lineages of similar size were more likely to yield false positives only when they included more than 4 rate shifts. We also compared the variance in lineage sizes between simulations for which we recovered the correct number of events against those for which we recovered too few events. For every possible number of lineages, we found significantly lower variance for simulations that were correctly assessed. Thus, we only see a slight effect of the lineage size on the occurrence of false positives, whereas high variance in lineage size significantly increases false negatives. This indicates on the one hand, a tendency to overestimate the number of shifts when lineages are comparable in size, and on the other hand, problems with Medusa for identifying diversification shifts specific to a low number of species, as showed in the first part.

**Figure 5 –.**
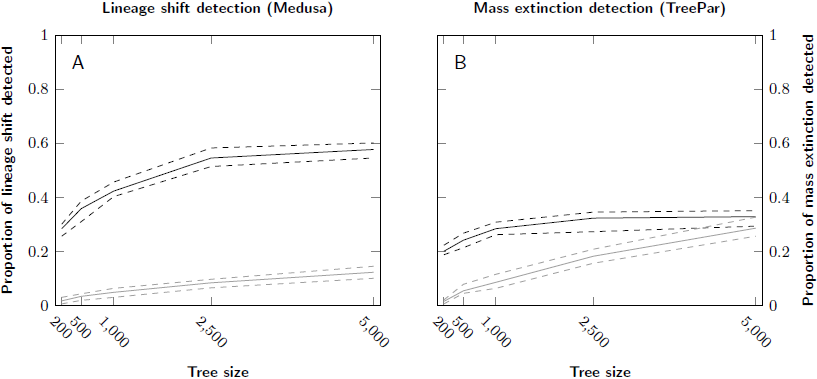
Influence of tree size on the detection of lineage shifts (A) and mass extinction events (B). Continuous lines correspond to median proportions of simulations and dotted lines correspond to 95% confidence interval, both based on resampling. Dark lines represent the proportion of simulations where the model with the correct number of events was the most favored, and light lines where a model with too many events was favored.

**Figure 6 –.**
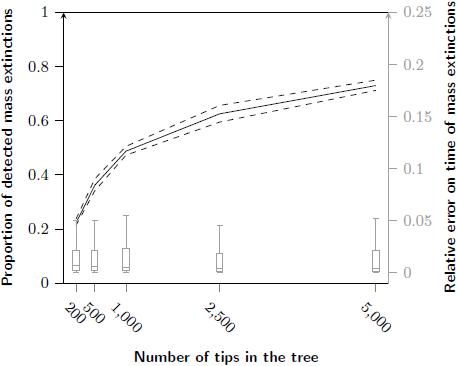
Influence of tree size on the detection of mass extinctions by TreePar. Line: proportion of detected mass extinctions; boxplots: distribution of the errors on their timing relative to the time of the first speciation event of the tree.

#### Impact of events violating the model

We tested the robustness of the methods by studying the behavior of (1) Medusa to detect rate shifts with an increasing number of mass extinctions, and (2) TreePar to detect mass extinction events with an increasing number of lineages shifts. The results obtained by Medusa were unaffected by the number of mass extinctions in the simulations (Fig 7). In contrast, an increase in the number of lineage shifts resulted in an increase of the proportion of false positives for TreePar (2% with no lineage shift vs. 20% with five; Fig. 7). However, the accuracy of the estimate of the timing of the event was not affected (Fig. 8). The number of lineage shifts had almost no impact on the probability of detecting a true mass extinction event, i.e. on false negatives.

**Figure 7 –.**
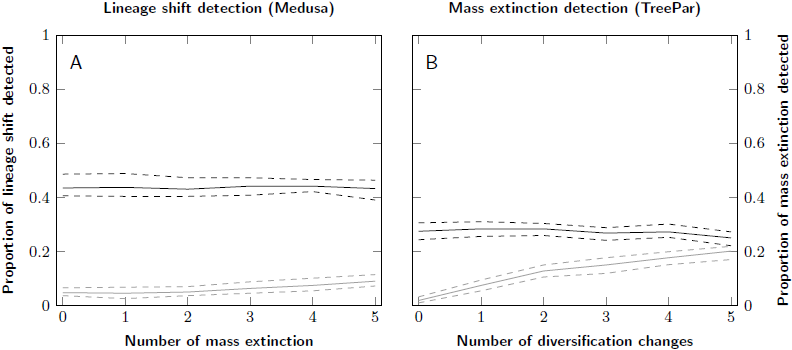
Influence of increasing model violations on the tests. A: Lineage shift detection against an increasing number of mass extinctions; B: Mass extinction event detection against an increasing number of lineage shifts. Dark lines: simulations where the correct number of events was found, light lines: simulations where too many events was favoured.

**Figure 8 –.**
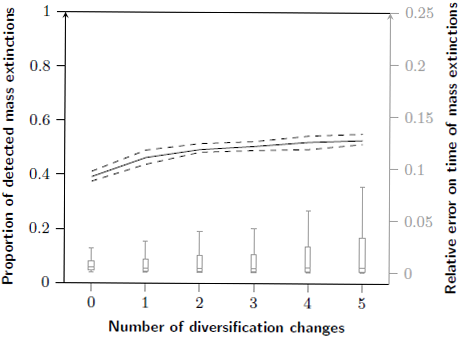
Influence of the number of lineage shifts in a simulation upon the detection of mass extinctions. Line: proportion of detected mass extinctions; boxplots: distribution of the errors on their timing relative to the time of the first speciation event of the tree.

#### Impact of patterns of extinction

The effect of different scenarios of mass extinction on the results of Medusa and TreePar are presented in Figure. 9. First, as expected, no effect of the extinction scenarios was observed on the detection of lineage rate shifts detected by Medusa (Fig. 9A). In contrast, the fair game and wanton destruction scenarios impacted the estimation made by TreePar. They produced, for comparable levels of detection, more false positives than the field of bullets which was used in the previous simulations (73% and 74% for fair and wanton against 58% for field of bullets, Fig. 9B). Irrespective of the type of mass extinction simulated, there are very few false negatives, i.e. at least one extinction was detected in almost every tree. The error on the timing of this event was kept under 5% of the root age. We also performed a search for global rate shifts in the case of wanton destruction (Fig. 9B, dashed background). Regarding this scenario, we also compared simulations where all lineages underwent an increase of diversification after the mass extinction event against those who underwent a decrease and observed no difference between the outcomes of the two frameworks. Even though the shifts were different between lineages (i.e., increase of diversification in some lineages, decrease in others), TreePar detected the period of this shift with more power than for the detection of the associated mass extinction (34% and 21% correctly assessed simulations, respectively). Overall, these results show that departure from the simplest model of mass extinction should not affect our ability to detect these events in phylogenetic trees (i.e. no increase in false negatives rate). But it should lead to an increase of false positive detections.

**Figure 9 –.**
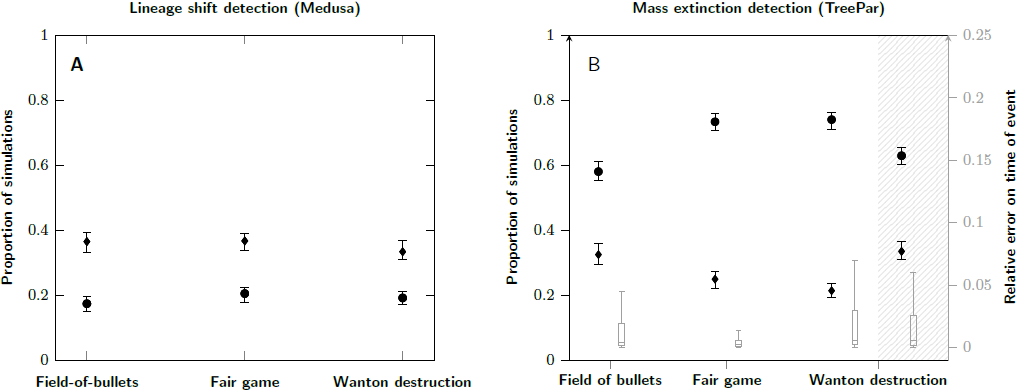
Influence of distinct extinction scenarios on Medusa and TreePar predictions. A: Medusa outcome; diamonds: proportion of simulations where the model with the correct number of events is chosen; circles: proportion of simulations where a model with too many events is chosen; there are less correctly assessed simulations for Medusa because of the high number of lineage shifts in these simulations (5). B: TreePar outcome and error on the timing of events: boxplots: error on the timing of the estimated extinction relative to the first speciation event; blank background: detection of mass extinctions; dashed background: detection of global rate shifts; other symbols as in A.

## CONCLUSION

Previous simulations involving mass extinctions and changes in macro-evolutionary rates have only focused on their effect on lineage through time plots (Crisp and Cook 2009). This lead to the identification of a possible mass extinction event in some plants lineages around 32 Mya, which was further suggested to be linked with changes in climate.

Recently, Hohna et al. Höhna (2013) developed a new algorithm to perform simulations with varying macro-evolutionary rates, allowing for mass extinction events. Other ongoing work aims at studying and simulating increasingly complex scenarios of diversification (Hartmann et al. 2010; Morlon et al. 2010). Condamine et al Condamine et al. (2013) have also discussed the possibility of modeling mass extinction events either as a single pulse of species loss or as an extended period of increase in the extinction rate. We have addressed here a different point, by investigating simultaneously different types of diversification processes, and assessing their effect on widely used method to estimate the dynamics of the diversification process.

The study of diversification rates has become a standard part of the analysis of large phylogenetic trees (Meredith et al. 2011; Jetz et al. 2012; Near et al. 2013), and recent efforts have also assessed the methods used when their assumptions are violated (Rabosky 2014). We have shown that departure from the assumption of consistency in rates across lineages causes a large increase in false positives when looking for mass extinction events. This can be problematic as we know that rate consistency rarely holds (Rabosky et al. 2007; Jetz et al. 2012; Barker et al. 2013), and casts doubts on our ability to reliably find such events using only phylogenetic trees. Nevertheless, an increasing number of disparities between lineages caused neither a decrease in the probability of detecting an event nor an increase in the error on its timing. As we observed the same pattern under more complex scenarios of extinction, the difficulty in detecting the K-Pg event in mammals is therefore probably not due to biases in the methods used. We might be limited by the power of TreePar to detect mass extinction events, although in simulations we reach 60% of true events detected for a tree size similar to that of mammals.

Recent efforts aim to reach a better agreement between paleontological and molecular data (Morlon et al. 2011), including looking for mass extinctions in molecular phylogenies. For instance, there is much debate on whether the K-Pg extinction event triggered the mammalian diversification (Bininda-Emonds et al. 2007; Meredith et al. 2011; Stadler 2011a; Dos Reis et al. 2012; O’Leary et al. 2013). The fossil record also indicates higher extinction rates of mammalians species around 65 Mya (Wilson 2005). In this work, we have shown that for phylogenetic trees similar in size to that of mammals (i.e. ca. 5000 species), the signal for mass extinctions was usually recovered in the tree, even though lineage discrepancies in macro-evolutionary rates had a tendency to yield more false positives. Thus, if the ancestor lineages of the extant mammal families did experience a mass extinction at the K-Pg boundary, we should theoretically be able to identify it using phylogenetic trees. The underlying assumption about the mass extinction made when using TreePar is that lineages are terminated randomly with a fixed *ρ* value everywhere in the tree, i.e. a field of bullets type of mass extinction. But other models of extinction seem to increase false positives but not false negatives, not explaining difficulties in finding a K-Pg signal in real phylogenetic trees.

Recent studies have used Markov processes to account for the effect of specific traits upon the probability of extinction of a species, thus extending models of mass extinction beyond the field of bullets (Faller and Steel 2012). Such models can be used for instance to estimate the loss of phylogenetic diversity after a mass extinction event (Lambert and Steel 2013). Our simulations can be seen as a special case of such models, where the trait influencing survival probabilities is the diversification value of the species. We have shown that more complex models of mass extinction cause more false positive detection than the simple field of bullets, as well as a decrease in the error for the fair game scenario.

Choosing a specific model of extinction (field of bullets, wanton destruction, fair game) might require incorporation of fossil information into the phylogeny, and thus the further development of methods capable of dealing with both molecular and fossil data.

## AUTHORS CONTRIBUTIONS

SL, MRR and NS designed the study, SL performed the simulations, SL, MRR and NS analysed the results and wrote the manuscript.

## ACKNOWLEDGEMENTS

This work was supported by the ProDoc grant number 134931 of the Swiss National Science Fondation; and État de Vaud. The computations were performed at the Vital-IT (http://www.vital-it.ch) Center for high-performance computing of the SIB Swiss Institute of Bioinformatics. We thank Tanja Stadler for helpful discussions.

